## Supplementary figures and images for "U2 snRNA structure is influenced by SF3A and SF3B proteins but not by SF3B inhibitors"

### Supporting Information SI1

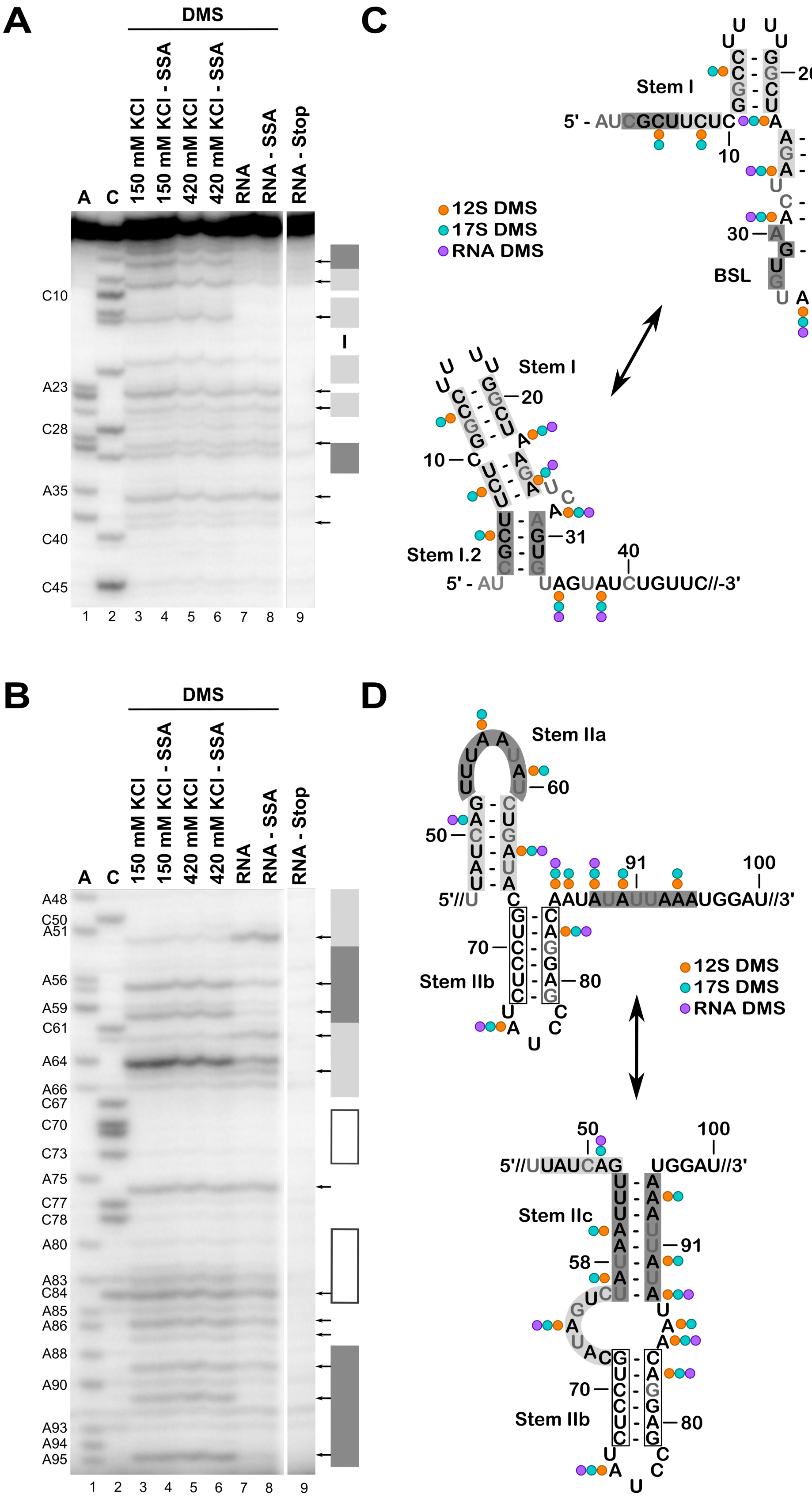

### Supporting Information SI2

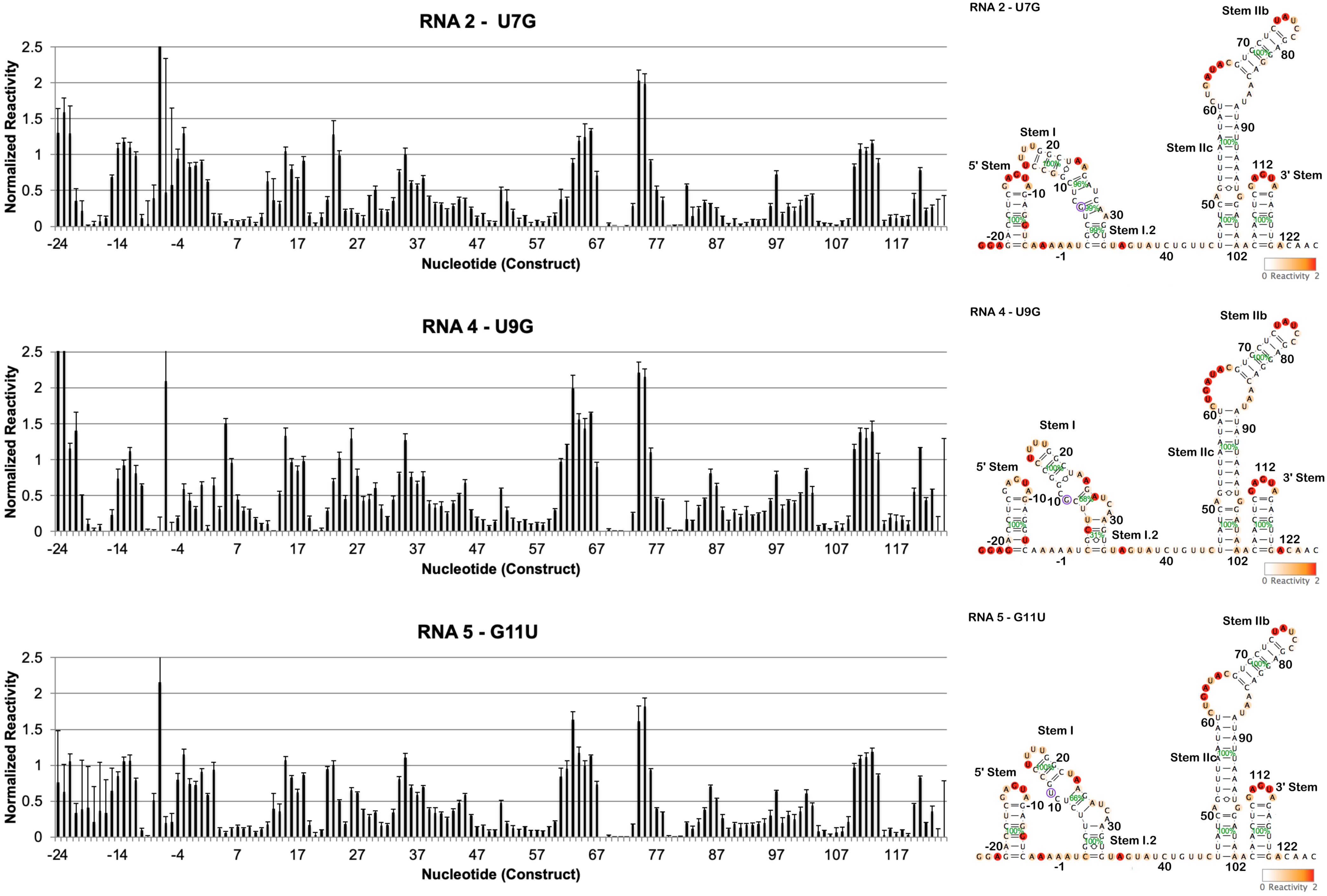

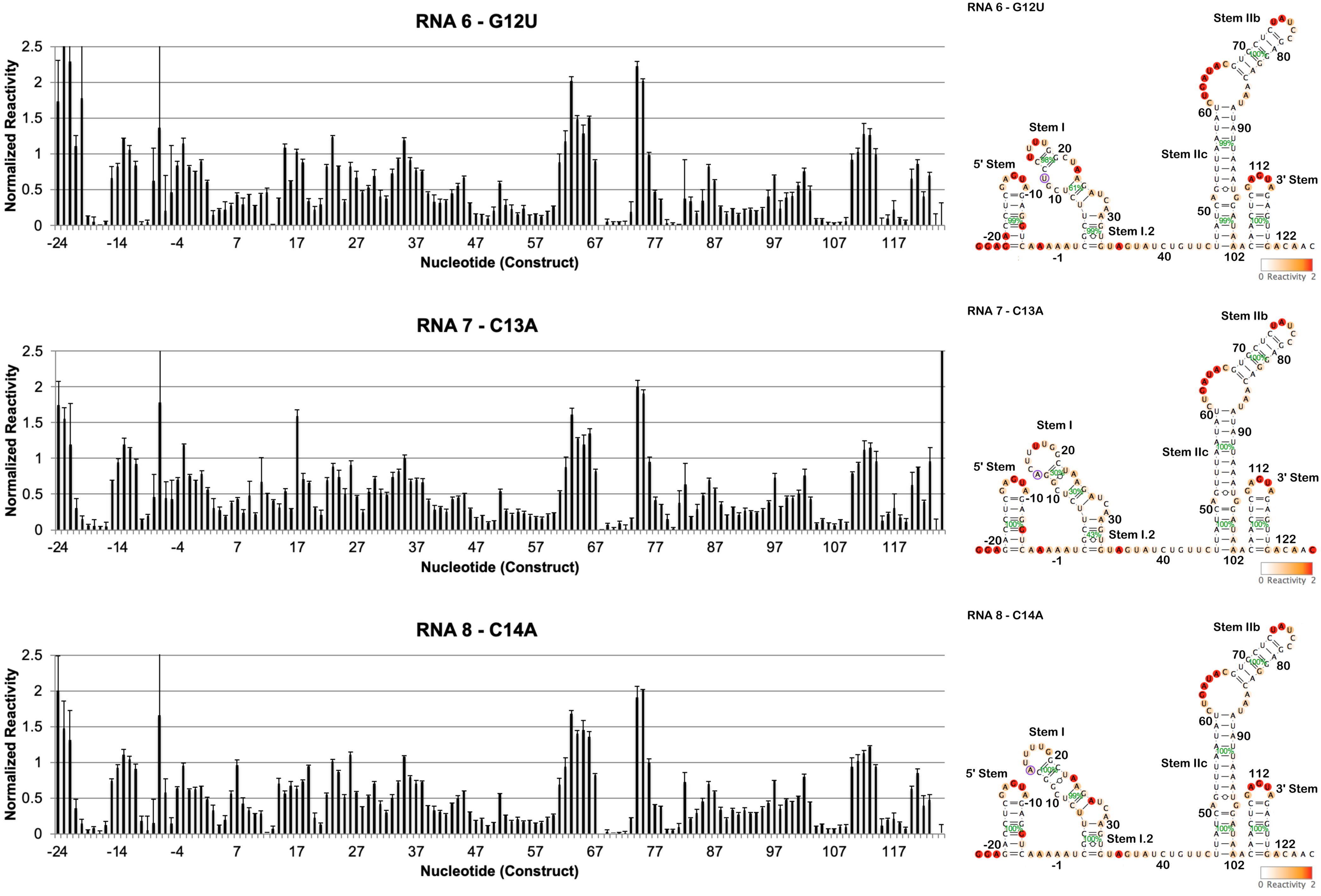

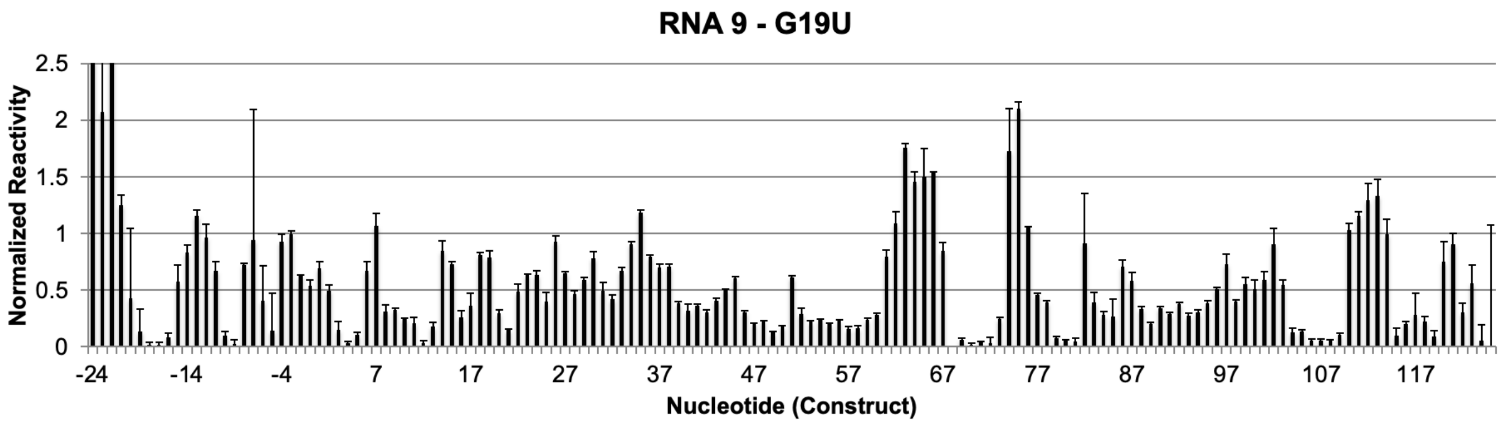

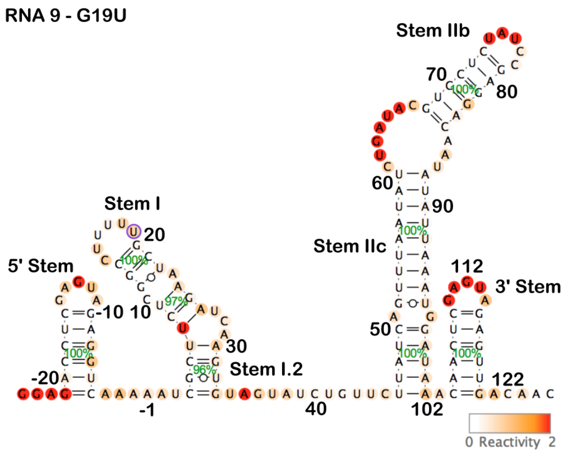

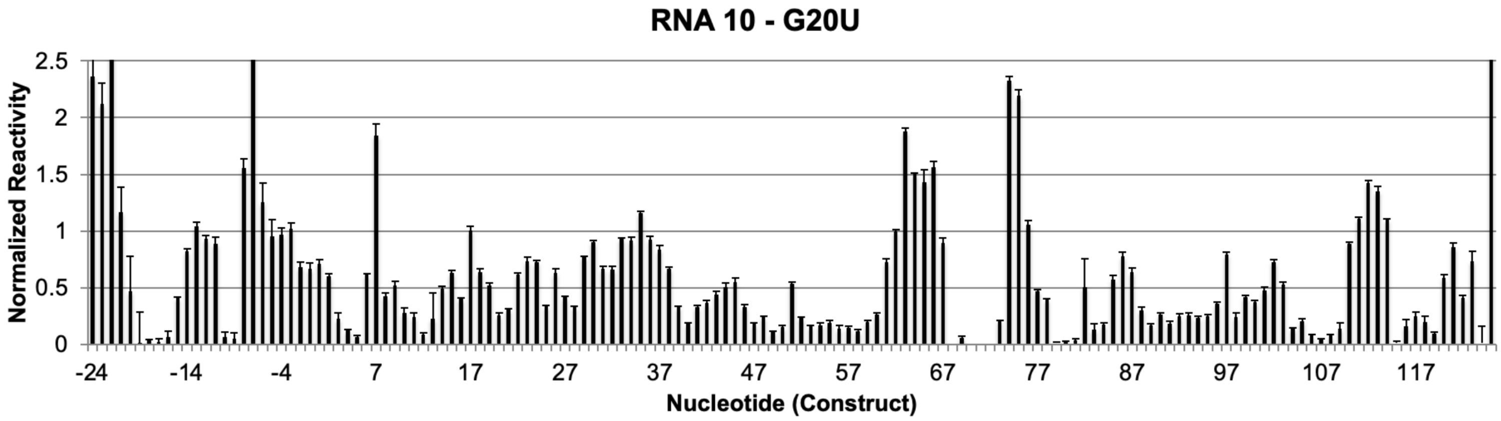

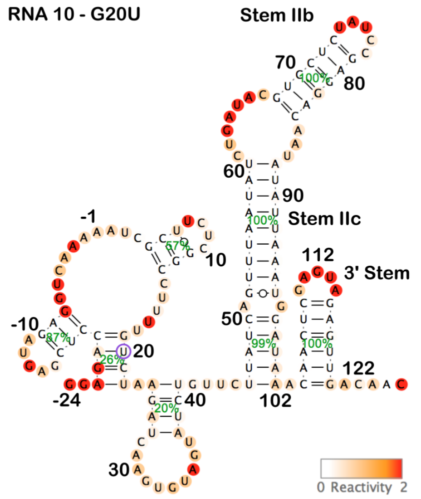

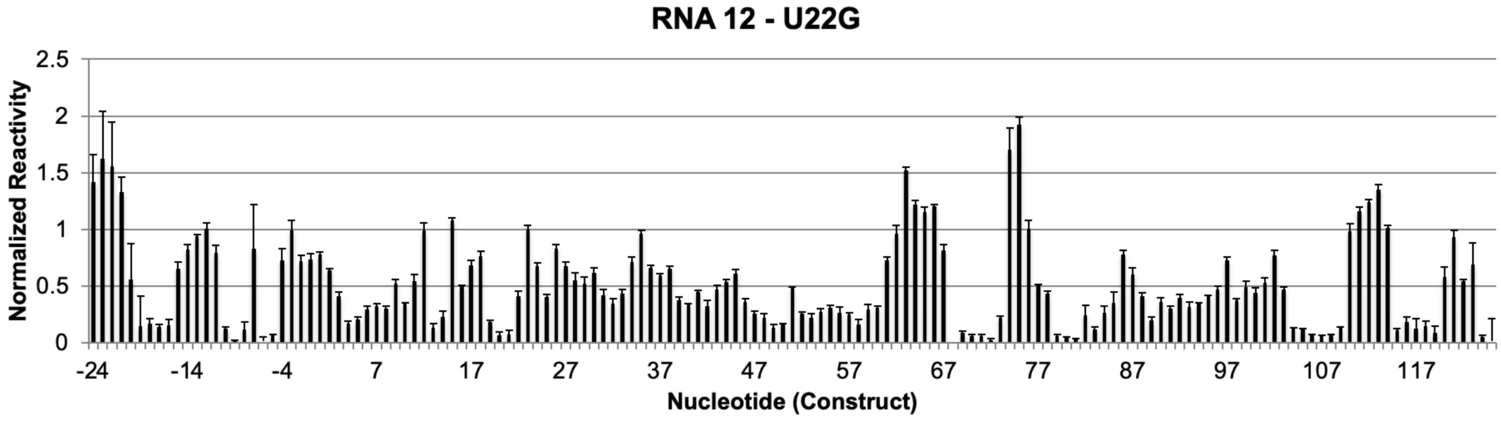

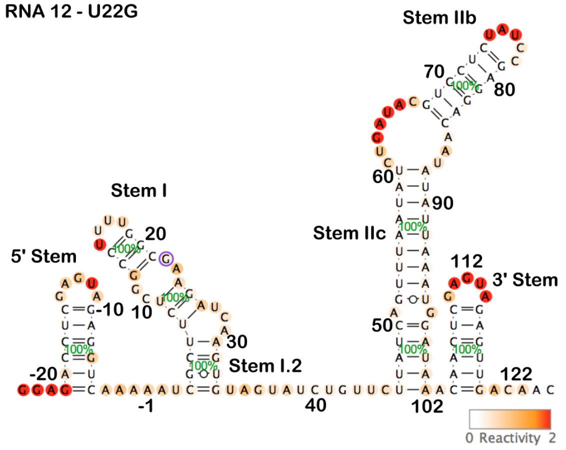

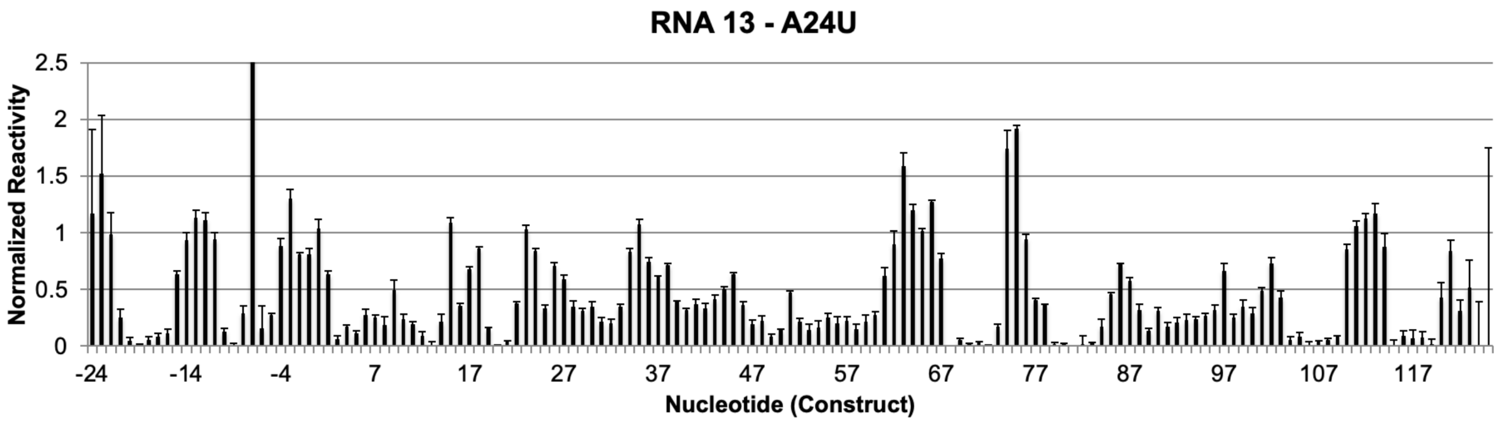

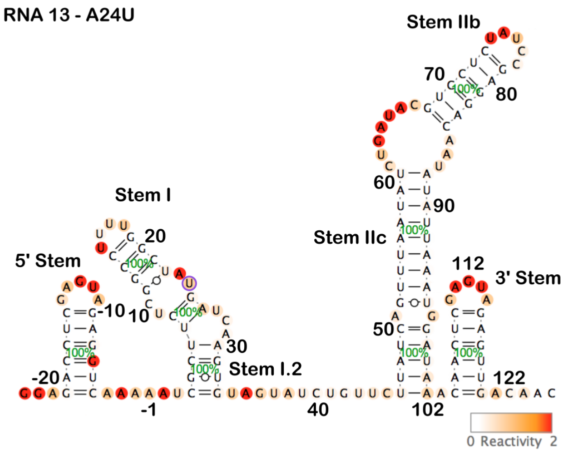

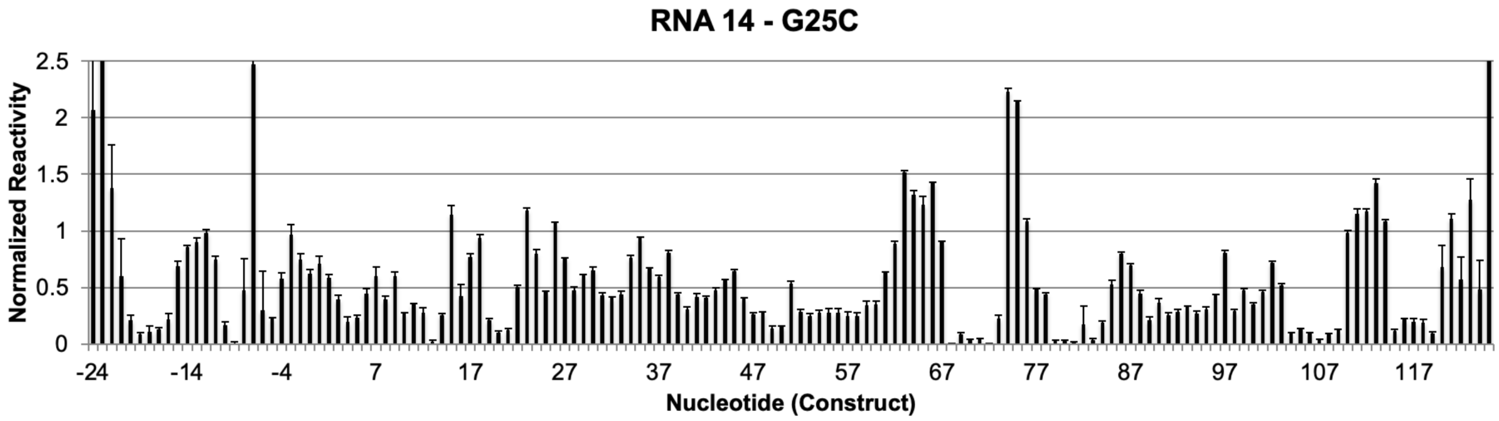

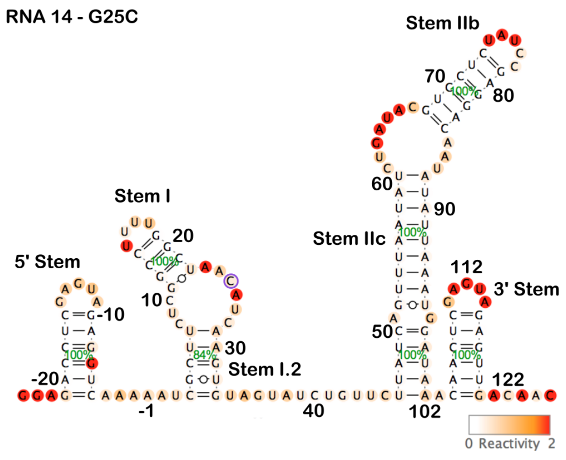
